# Ligament injury in adult zebrafish triggers ECM remodeling and cell dedifferentiation for scar-free regeneration

**DOI:** 10.1101/2023.02.03.527039

**Authors:** Troy Anderson, Julia Mo, Ernesto Gagarin, Desmarie Sherwood, Maria Blumenkrantz, Eric Mao, Gianna Leon, Hung-Jhen Chen, Kuo-Chang Tseng, Peter Fabian, J. Gage Crump, Joanna Smeeton

**Affiliations:** Columbia Stem Cell Initiative, Department of Rehabilitation and Regenerative Medicine, and Department of Genetics and Development, Columbia University Irving Medical Center, Columbia University, New York, NY 10032, USA; Department of Biological Sciences, Columbia College, Columbia University NY 10027, USA; Packer Collegiate Institute, New York, NY 11201, USA; Department of Stem Cell Biology and Regenerative Medicine, Keck School of Medicine, University of Southern California, Los Angeles, CA 90033, USA

**Keywords:** ligament, joint, zebrafish, regeneration, single-cell RNA sequencing, legumain, dedifferentiation

## Abstract

After traumatic injury, healing of mammalian ligaments is typically associated with fibrotic scarring as opposed to scar-free regeneration. In contrast, here we show that the ligament supporting the jaw joint of adult zebrafish is capable of rapid and complete scar-free healing. Following surgical transection of the jaw joint ligament, we observe breakdown of ligament tissue adjacent to the cut sites, expansion of mesenchymal tissue within the wound site, and then remodeling of extracellular matrix (ECM) to a normal ligament morphology. Lineage tracing of mature ligamentocytes following transection shows that they dedifferentiate, undergo cell cycle re-entry, and contribute to the regenerated ligament. Single-cell RNA sequencing of the regenerating ligament reveals dynamic expression of ECM genes in neural-crest-derived mesenchymal cells, as well as diverse immune cells expressing the endopeptidase-encoding gene *legumain*. Analysis of *legumain* mutant zebrafish shows a requirement for early ECM remodeling and efficient ligament regeneration. Our study establishes a new model of adult scar-free ligament regeneration and highlights roles of immune-mesenchyme cross-talk in ECM remodeling that initiates regeneration.

**Highlights:**

- Rapid regeneration of the jaw joint ligament in adult zebrafish
- Dedifferentiation of mature ligamentocytes contributes to regeneration
- scRNAseq reveals dynamic ECM remodeling and immune activation during regeneration
- Requirement of Legumain for ECM remodeling and ligament healing

## Introduction

Ligament injuries typically heal with biomechanically inferior scar tissue that compromises joint stability and increases the risk of developing painful and debilitating osteoarthritis^1,2^. Healed ligaments fail to recapitulate the highly organized structure of the uninjured tissue. Foundational research using non-regenerative models has highlighted that the failure to regenerate ligaments without scarring is common among mammals^3–5^. Early studies investigating repair of the medial collateral ligament (MCL) following rupture showed that the healed ligament undergoes more deformation under the same load, coincident with disorganization of the collagen fibers at the wound site^5,6^. The first 10 days of ligament healing in mammals are characterized by an invasion of fibroblasts, vasculature, and pro-inflammatory immune cells along the ligament^6,7^. While ligament repair continues for 14 weeks, an incomplete remodeling of both the cellular and extracellular matrix (ECM) composition occurs, ending with a fibrotic scar bridging the injury. Similarly, in human and murine tendon and ligament injuries, the stubs are bridged by α-Smooth Muscle Actin-positive scarring myofibroblasts^8–11^. Notably, these myofibroblasts do not express tendon and ligament marker *Scleraxis* (*scx*), indicating that they are not true ligamentocytes^8^. Existing studies on adult connective tissue injury highlight the importance of the reestablishment of cell fate and ligament ECM through regeneration.

Recent advances in tendon injury modeling showed that neonatal mice are capable of scar-free regeneration after Achilles tendon transection^8^. In this neonatal repair context, *Scx*-positive tenocytes retain their developmental potency to proliferate, migrate to the wound site, and heal without forming a fibrotic scar^8^. Further studies on developmental regenerative potential have been performed using juvenile zebrafish, which described that BMP-dependent activation of perichondrial progenitors is necessary for tendon regeneration after tendon cell ablation injury^12^. However, while these models interrogate scar-free tendon regeneration in the context of development, there are currently no models to study scar-free ligament regeneration in mature adult ligaments. To address this need, we utilized the highly regenerative zebrafish as they are a well-established model for heart, spinal cord, and tail fin regeneration^13–15^ and moreover possess regenerative synovial joints supported by ligaments^16–18^.

In the course of establishing a post-traumatic osteoarthritis model, we had observed rapid regeneration of the craniofacial interopercular-mandibular (IOM) ligament, which supports the zebrafish jaw joint, following complete transection in one-year-old fish^18^. Here we characterize the cellular and molecular events underlying the three overlapping phases of ligament healing: inflammation, proliferation, and remodeling. We also used single-cell RNA sequencing (scRNAseq) to characterize the dynamics of cell populations during regeneration and identified *legumain* (*lgmn*) as a gene highly enriched in macrophages that invade the injury site. Legumain is a member of the C13 family of cysteine proteases that regulate ECM factors such as cathepsins, fibronectin1 (Fn1), and matrix metalloprotease-2 (MMP-2)^19,20^. By generating a zebrafish mutant for *lgmn*, we uncovered an essential role in regulating ECM remodeling and scar-free ligament regeneration. Our findings highlight the important role of immune-mediated ECM remodeling for successful scar-free regeneration.

## Results

### Complete and scar-free regeneration of the IOM ligament following transection

The IOM ligament connects the interopercular bone to the retroarticular bone of the mandible to control movement of the zebrafish jaw joint. Following complete transection of the IOM ligament in adult zebrafish, a time-course of histological analysis revealed dynamic morphological changes during ligament regeneration (Fig. 1a-h). The uninjured ligament contains ligamentocytes with elongated nuclear morphology that are surrounded by a cell-sparse ECM (Fig. 1a,b). Within 1 day post-ligament transection (dplt), a robust inflammatory response is seen that begins to resolve by 3 dplt. From 1 to 3 dplt, we observe breakdown of ligament ECM and rounding of ligament-embedded nuclei (Fig. 1c-d). Beginning at 3 dplt and continuing through 7 dplt, abundant mesenchymal cells are observed in the gap between the injured ligament stubs (Fig. 1d-e). By 14 dplt, cells bridging the RA and IOP bones display an elongated morphology in the direction of force but at a high cell density compared to native uninjured ligaments (Fig. 1f). From 21 to 28 dplt, the hypercellularity is resolved and the regenerated ligament resembles the cell-sparse uninjured ligament (Fig. 1g-h). These data point to a transient inflammatory response, ligament stub ECM breakdown, followed by establishment of a population of mesenchymal cells bridging the wound site that are remodeled into the regenerated ligament.

**Figure 1.**
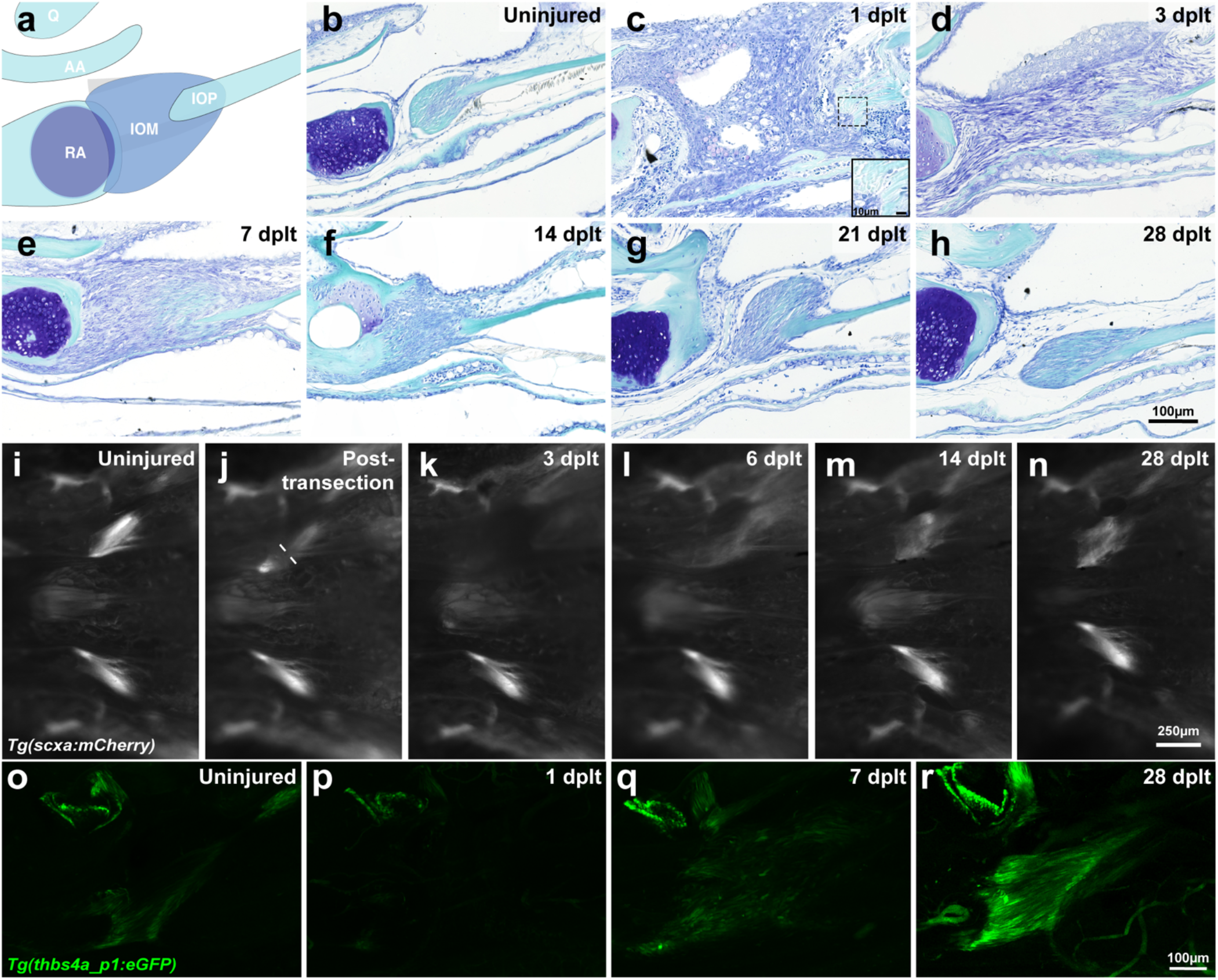
Adult zebrafish ligament transection injury and regeneration. **(a)** Schematic of the articulating bones at the zebrafish jaw joint and connections of the interopercular-mandibular (IOM) ligament. **(b-h)** Toluidine blue staining of adult zebrafish IOM ligament in uninjured (b, n=16), 1 day post ligament transection (dplt, c, n=3), 3 dplt (d, n=5), 7 dplt (e, n=6), 14 dplt (f, n=16), 21 dplt (g, n=3), and 28 dplt (h, n=16). **(i-n)** Repeated imaging of the ventral view of *scxa:mCherry* expression in IOM ligament before injury (i, n=9), immediately after injury (j, n=3), 3 dplt (k, n=9), 6 dplt (l, n=3), 14 dplt (m, n=9), and 28 dplt (n, n=3). **(o-r)** confocal microscopy imaging of tissue cleared *thbs4a_p1:e1b-GFP* expression in IOM ligament uninjured control (o, n=6), 1 dplt (p, n=5), 7 dplt (q, n=6), and 28 dplt (r, n=3). AA, anguloarticular bone; IOM, interopercular-mandibular ligament; IOP, interopercular bone; Q, quadrate bone; RA, retroarticular bone. Scale bars= 100 μm (b-h), 250 μm (i-n).

To understand the activity of ligamentocytes during regeneration, we labeled these cells with a *scxa:mCherry* transgenic line. In these fish, mCherry expression is driven by the regulatory region surrounding the *scxa* gene, which encodes a transcription factor highly expressed in ligament and tendon cells^21,22^. In uninjured adults, live imaging revealed *scxa*:mCherry fluorescence in each of the bilateral IOM ligaments (Fig. 1i). Immediately following the transection, we observed decreased *scxa*:mCherry fluorescence only on the injured side, with mCherry fluorescence nearly completely absent by 3 dplt. We then saw reappearance of *scxa*:mCherry fluorescence by 6 and 14 dplt, gradually returning to near-normal levels by 28 dplt (Fig. 1j-m). This result suggests while ligament stub-resident ligamentocytes were observed in histological analysis, these mature cells show a dramatic loss of *scxa*:mCherry expression adjacent to the transection site.

Expression of *scxa* is known to be broader than just tenocytes and ligamentocytes, including early mesenchyme and fibroblasts^23^. We therefore sought to confirm our results with a transgenic line marking mature ligamentocytes more specifically. Analysis of our published single-cell assay for transposase-accessible chromatin (scATAC) sequencing data of zebrafish cranial neural crest-derived cells^24^ identified a 635 bp region within intron 13 of the *thbs4a* gene that was selectively accessible in the tendon and ligament cluster (Fig. S1a). We then used this sequence to generate transgenic *thbs4a_p1*:eGFP zebrafish. In contrast to expression of *scxa*:mCherry throughout tendons and ligaments, *thbs4a_p1*:eGFP expression was highly specific for the IOM ligament, with the exception of a small amount of superficial cartilage expression in the jaw joint (Fig. 1o; Fig. S1b). Following IOM ligament transection, we observed disappearance of *thbs4a_p1*:eGFP expression in spared ligament domains flanking the cut site by 1 dplt, reappearance by 7 dplt, and remodeling to a normal ligament morphology by 28 dplt (Fig. 1o-r, Videos S1-4). In particular, the use of a CUBIC tissue clearing technique revealed that ligamentocytes had re-established alignment in the direction of force at 28 dplt, in contrast to the disorganized alignment seen at 7 dplt. The loss of expression of two independent reporters in ligamentocytes flanking the transection site strongly suggest that remaining ligamentocytes dedifferentiate early in the regenerative process. Further, re-expression of these reporters later in the regenerative process show re-activation of the developmental gene regulatory program for ligamentocytes during adult regeneration.

### Mature ligamentocytes contribute to the regenerated ligament

Next, we sought to define the source of ligamentocytes during regeneration. During development, the IOM ligament and other components of the craniofacial skeleton are neural crest-derived. We therefore tested whether the regenerative mesenchyme and regenerated ligament are similarly derived from neural crest-derived cells. To do so, we permanently labeled neural crest lineage cells in *Sox10:Cre;actb2:loxP-BFP-STOP-loxP-DsRed* fish, in which *Sox10*-driven Cre recombinase switches blue fluorescent protein (BFP) to DsRed fluorescent protein (Fig. 2a). Repeated live-imaging through the course of ligament healing revealed the regenerative mesenchyme to be DsRed+ at 3, 7, and 14 dplt, and the regenerated ligament DsRed+ at 14 and 28 dplt. Thus, similar to their development, both the early bridging regenerative mesenchyme and regenerated ligament are produced by neural crest-derived cells.

**Figure 2.**
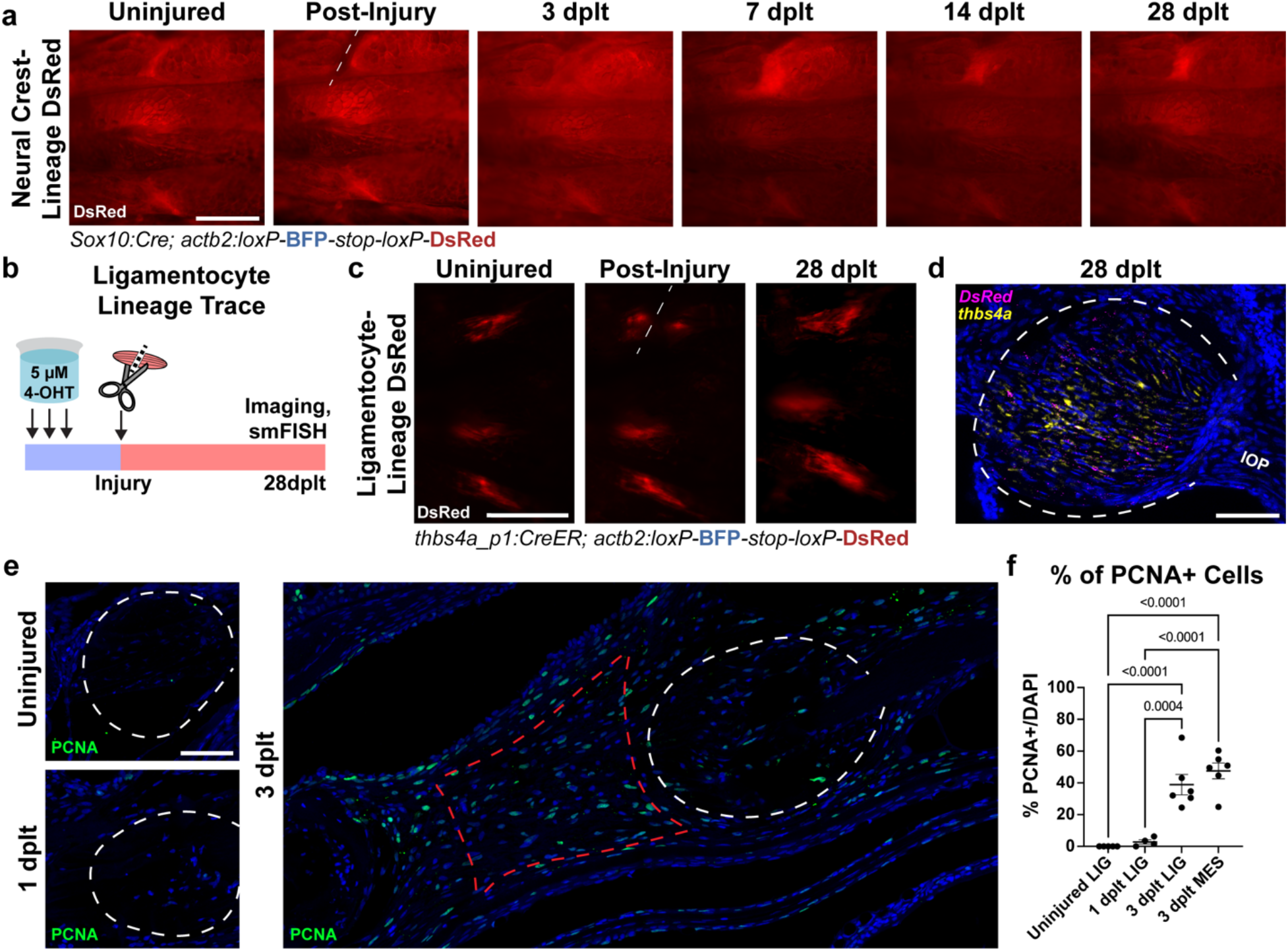
Lineages contributing to ligament regeneration following transection injury. **(a)** Lineage trace of DsRed-labeled neural crest-derived cells using sox10:Cre with actb2:loxP-BFP-STOP-loxP-DsRed through ligament regeneration at 0 hours post-ligament transection (Injury) (n=5). **(b)** Schematic of the ligamentocyte lineage tracing experiment. 5-month old fish were treated with 5 μM 4-hydroxytamoxifen (4-OHT) overnight three times and screened for conversion, then imaged before ligament transection, immediately after transection (Injury), and at 28 dplt. **(c)** Live imaging from a ventral view of one fish before, immediately following, and 28 days post ligament transection (Cut site represented by the dashed white line) (n=8). **(d)** RNAscope single-molecule in situ hybridization of the regenerated ligament illustrating co-localization of DsRed (magenta) transcripts with *thbs4a* (yellow) transcripts, and nuclei stained with DAPI in blue (n=5). **(e)** PCNA immunofluorescence (green) and DAPI nuclear stain (blue) in uninjured, 1 dplt, and 3 dplt tissue sections. Ligament stub domains outlined in white dashed line, 3 dplt regenerative mesenchyme domain in red dashed line. **(f)** Quantification of percentage of PCNA+ nuclei in (e), showing a significant increase in proliferation at 3 dplt (n=4-6 per time point) relative to uninjured and 1 dplt ligaments. Scale bars= 500 μm (a,c), 50 μm (d,e).

We next sought to understand what subset of neural crest-derived cells might contribute to the regenerated ligament. Given our finding that mature ligament cells undergo rapid dedifferentiation in response to ligament transection, we tested their potential long-term contribution to the regenerated ligament. To permanently label ligamentocytes prior to injury, we used the *thbs4a_p1* ligament-specific enhancer to drive expression of 4-hydroxy tamoxifen (4-OHT) inducible CreERT2 (*thbs4a_p1:CreERT2*) in combination with the ubiquitously-expressed *actb2:loxP-BFP-STOP-loxP-DsRed* switch reporter. Treatment of adult double transgenic fish with 3 doses of 4-OHT resulted in DsRed fluorescence throughout the IOM ligaments (Fig. 2b,c). Following ligament transection, repeated live-imaging revealed the regenerated ligament to be DsRed+ at 28 dplt, with DsRed expression co-localizing with endogenous expression of *thbs4a* in ligamentocytes (Fig. 2c,d). In addition, we observed extensive proliferation, based on immunofluorescence staining for Proliferating Cell Nuclear Antigen (PCNA), within both the regenerative mesenchyme and spared ligament tissue adjacent to the transection site. Compared to only minimal PCNA+ cells prior to injury and at 1 dplt, 39% of cells within the ligament stub domain and 47.5% cells within the regenerative mesenchyme domain were PCNA+ at 3 dplt (Fig. 2e-f, n=4-6 per sample). These data support typically quiescent ligamentocytes re-entering the cell cycle upon injury and contributing to the regenerated ligament.

### Single-cell RNA sequencing reveals dynamic mesenchymal populations during ligament regeneration

To gain insights into the cellular and molecular dynamics underlying ligament regeneration, we performed droplet-based single-cell RNA sequencing (scRNAseq) on micro-dissected IOM ligaments, including adjacent joint and mesenchymal tissues, before and during early stages of regeneration. We used FACS in *Sox10:Cre;actb2:loxP-BFP-STOP-loxP-DsRed* fish to enrich for neural crest-derived cells, yet sequenced both DsRed+ neural crest-derived and BFP+ non-neural-crest-derived populations to capture the entire diversity of cell types during regeneration. DsRed+ and BFP+ populations were sequenced independently from uninjured controls, and from fish at 1 and 3 days after IOM ligament transection or undergoing sham surgery to cut the skin but not the ligament (Fig. 3a). After pre-processing and filtering, we obtained 9,451 cells (1618 DsRed+, 7833 BFP+) with a median 766 genes per cell (Table S1).

**Figure 3.**
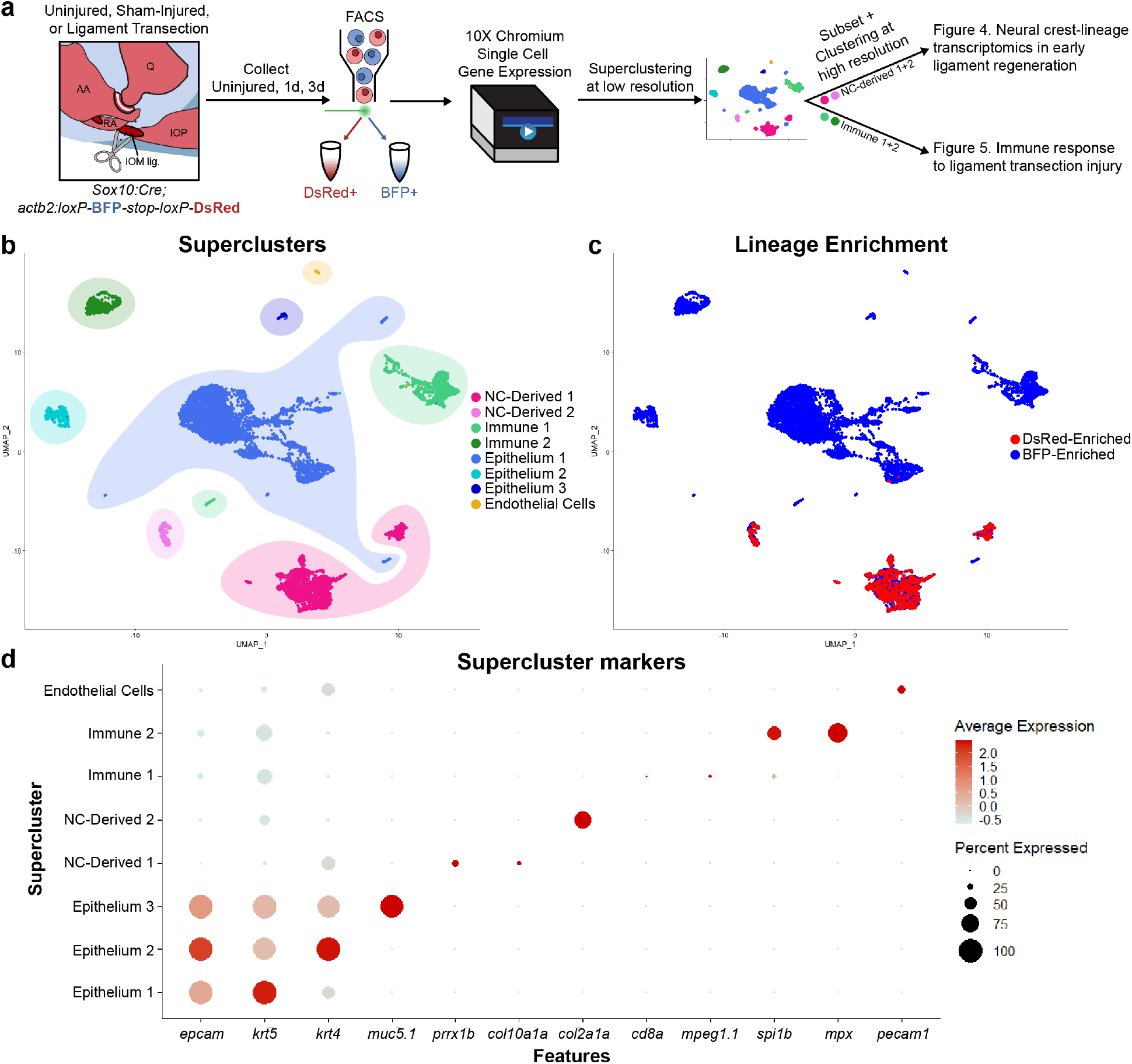
Single-cell transcriptomics of joint tissue through early ligament regeneration. **(a)** Schematic detailing the generation and analysis of single-cell RNA sequencing data. Jaw joints from uninjured, 1 day post-SHAM (nicking of the epithelium above the ligament) surgery, 3 days post-SHAM surgery, 1 day post-ligament transection, and 3 days post-ligament transection were dissociated and sorted for DsRed (neural crest-derived) or BFP (non-neural crest-derived). **(b)** UMAP of low-resolution superclusters for merged samples. Superclusters include 2 neural crest-derived clusters (NC-Derived), 2 immune clusters, 3 epithelial clusters, and 1 endothelial cluster. **(c)** UMAP colored for neural crest-lineage enrichment. Cells from FACS-sorted DsRed+ neural crest-derived cells colored in red, cells from sorted BFP+ non-neural crest-derived cells colored in blue. **(d)** DotPlot detailing expression of marker genes for each supercluster.

Unbiased, low-resolution clustering categorized cells into eight broad superclusters representing immune, epithelial, neural-crest-derived (NC-derived), and endothelial cells (Fig. 3b,c). Epithelial superclusters included basal and intermediate keratinocytes (Epithelium 1, *krt5*+), superficial cells (Epithelium 2, *krt4+*), and mucosal cells (Epithelium 3, *muc5.1+*) (Fig. S2). Immune 1 supercluster includes lymphoid and myeloid cells, such as T-cells (*cd8a+, lck+*) and macrophages (*mpeg1.1+, mfap4+*), and Immune 2 cluster consisted primarily of neutrophils (*mpx+, lyz+*). We also detected an endothelial cluster (*pecam1+, kdrl+*). Two superclusters, enriched for DsRed+ cells (Fig. 3c), represent neural-crest derived mesenchymal populations marked by either the broad mesenchymal gene *prrx1b* (NC-Derived 1) or the chondrocyte genes *col2a1a* and *mia* (NC-Derived 2) (Fig. 3d).

Re-clustering of the two neural crest-derived superclusters revealed clusters of osteoblasts (*ifitm5*+), chondrocytes (*col2a1a*+), superficial chondrocytes and synoviocytes (*prg4b*+), ligamentocytes (*scxa*+, *thbs4a*+), two types of dermal fibroblasts (*pah*+), periosteal cells (*mmel1*+), three types of stromal cells (defined by either *cxcl12a, coch*, or *col5a3a*), and glial cells (*mbpa+*)^18,24^ (Fig. 4a-c, Fig. S2, Table S2). In addition, we identified two injury-specific clusters with stage-specific cell composition: 1 dplt regenerative mesenchyme (66% 1 dplt cells) and 3 dplt regenerative mesenchyme (82% 3 dplt cells). These regenerative mesenchyme clusters were characterized by high expression of *collagen 12 alpha 1a* (*col12a1a*) and *fibronectin 1a* (*fn1a*) (Fig. 4c; Fig. S2). In situ hybridization shows that *col12a1a* is highly expressed in the mesenchymal cells bridging the ligament stubs by 3 dplt (Fig. 4d). Further, combinatorial in situ hybridizations show that a subset of *col12a1a+* regenerative mesenchyme cells co-express the early ligamentocyte marker *scxa*, suggesting early initiation of the ligament program by 3 dplt (Fig. 4d). In contrast, *col12a1a+* cells do not co-express *alpha-smooth muscle actin 2* (*acta2*), a marker of pro-scarring fibroblasts in mammals. These data are consistent with regenerative mesenchyme cells producing new ligamentocytes as opposed to scar tissue during regeneration.

**Figure 4.**
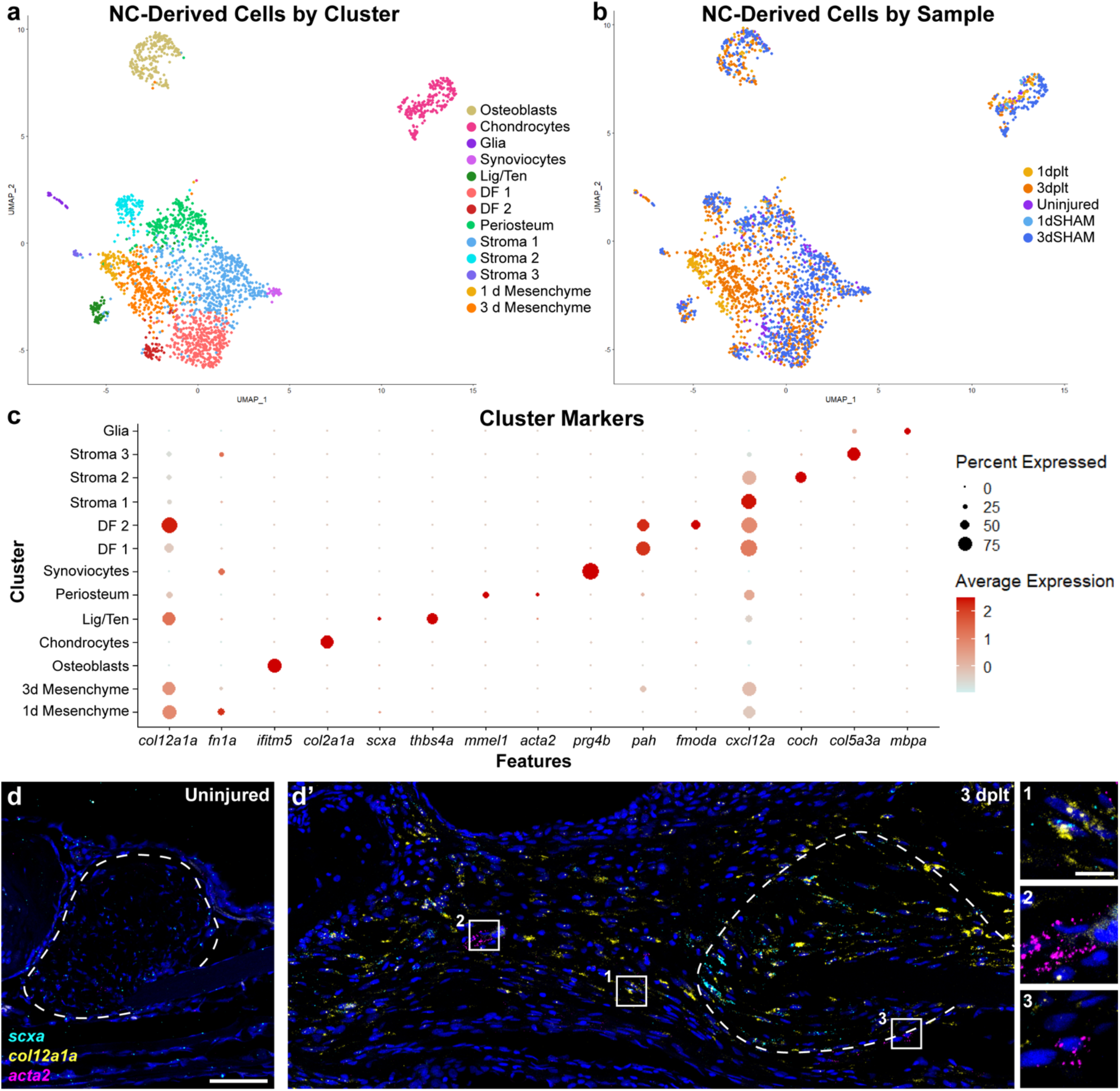
Neural crest-lineage single-cell transcriptomics in early ligament regeneration. (**a,b**) UMAP of single-cell RNA sequencing data of neural crest-derived cells from early ligament regeneration colored by (a) cluster or (b) sample. (**c**) DotPlot of marker genes used to identify each cluster. (**d**) RNAscope single-molecule fluorescent *in situ* hybridization (smFISH) for *scxa* (cyan), *col12a1a* (yellow), and *acta2* (magenta) in an uninjured ligament. (d’) smFISH for *scxa, col12a1a*, and *acta2* expression at 3 days post-ligament injury. Insets highlighting *scxa+ col12a1a+* regenerative mesenchyme (inset 1, n=6), *acta2+* perivascular cells (inset 2, n=5), and *acta2-low* mesenchyme near the ligament stub (inset 3, n=5). Scale bars = 50 μm (d, d’), 10 μm (insets in d’), ligament stub outlined in white dashed line.

### Single-cell profiling the immune response to ligament injury

Re-clustering of the two immune cell superclusters revealed 11 clusters: two types of neutrophils (*mpx*+), macrophages (*mpeg1.1+/mfap4+*), dendritic cells (*mpeg1.1 mfap4-*), two types of T helper cells (*il21r.2+* or *il4*+), proliferative cells (*mki67+*), and four types of lymphocytes (*lck*+) including two T cell-containing clusters (*cd8a+*), NK cells (*s1pr5a*+)^25^, and activated NK cells (*ifng1+/gzmk+*) (Fig. 5a-c, Fig. S3). The Neutrophil 1 cluster consisted predominantly of cells from the 1 dplt sample (82% of cells), indicative of an acute inflammatory phase shortly after injury.

**Figure 5.**
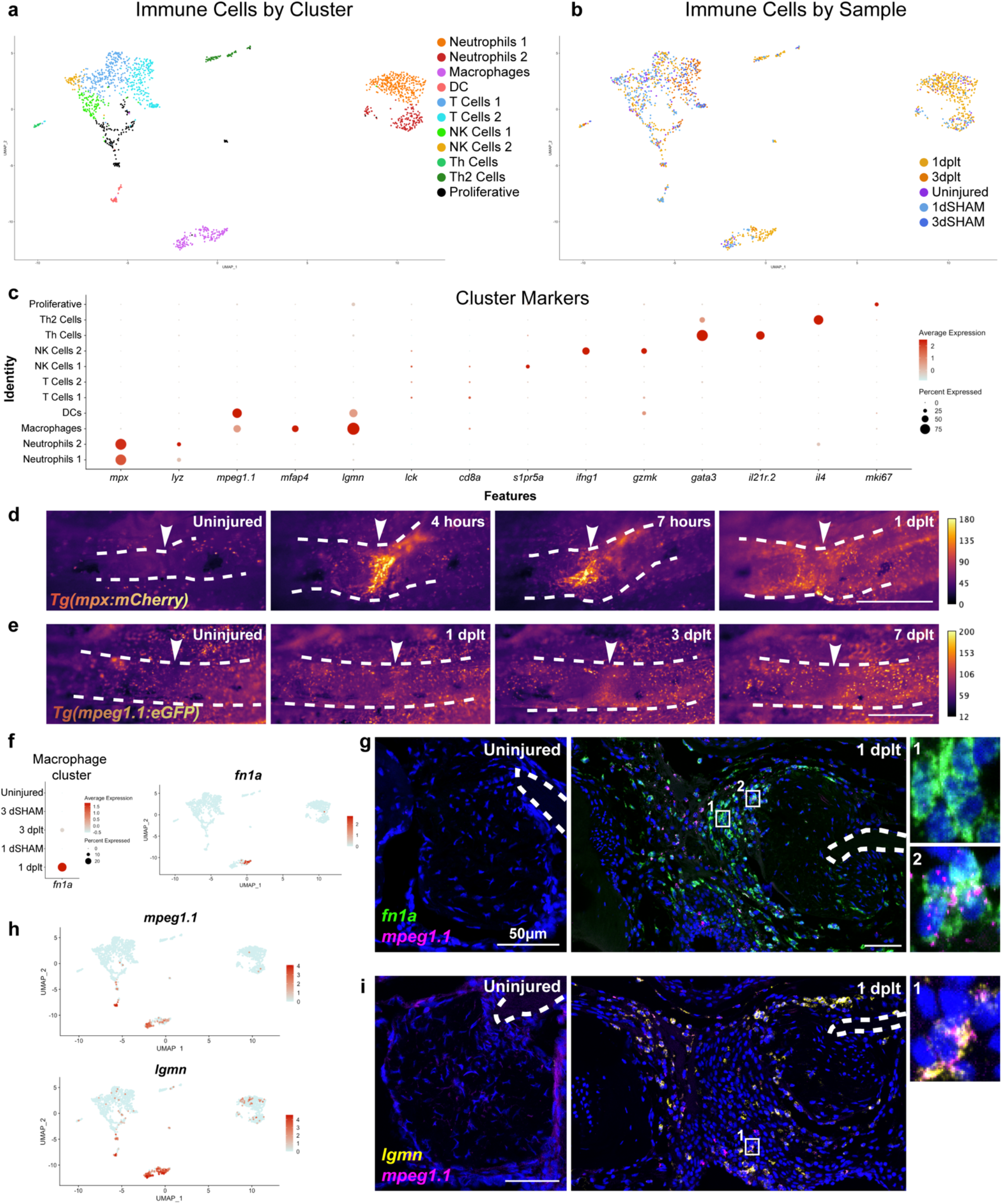
Immune response to ligament transection injury. **(a,b)** UMAPs of single cell RNA sequencing datasets color-coded by cluster (a) and injury sample (b). **(c)** Dot plot showing marker gene expression for each immune cluster. **(d,e)** Repeated live imaging of neutrophils (d; *mpx*:mCherry) (n=5) and macrophages (e; *mpeg1.1*:eGFP+) (n=6) at the injury site (arrowheads) during the first week of regeneration with color coded by signal intensity. (**f**) DotPlot demonstrating *fn1a* expression in macrophage cluster cells split by Sample and FeaturePlot for 1 dplt-macrophage enriched gene *fn1a*. (**g**) Representative images of smFISH staining using *fn1a* (green) and *mpeg1.1* (magenta) probes in IOM ligaments from uninjured and 1 dplt samples (n=3). (**h**) Feature plots for macrophage marker *mpeg1.1* and macrophage-enriched gene *lgmn*. (**i**) smFISH for *lgmn* (yellow) and *mpeg1.1* (magenta) probes in IOM ligaments from uninjured and 1 dplt samples (n=3). White dashed lines outline the IOP bone and inset positions are denoted by the numbers 1-2 (g, i). Scale bars = 500 μm (d,e), 50 μm (g,i).

Repeated live imaging of individual fish expressing the transgenic neutrophil reporter *mpx*:mCherry revealed homing of *mpx*+ neutrophils to the injury site within 4 hours, with neutrophil accumulation beginning to resolve by 1 dplt (Fig. 5d, white arrowheads). In contrast, repeated imaging of fish transgenic for the macrophage and dendritic cell reporter *mpeg1.1:eGFP* showed *mpeg1.1+* immune cells homing to the transection site at 1 dplt and peaking by 3 dplt, with skin-associated macrophages apparent similarly in uninjured and transected fish (Fig. 5e, white arrowheads).

Within the *mpeg1.1+/mfap4+* macrophage cluster, we noted a subpopulation of macrophages with enriched composition of 1 dplt cells and selective expression of *fn1a* (Fig. 5f). In situ hybridization at 1 dplt revealed *mpeg1.1+/fn1a+* macrophages close to the transection site; adjacent *mpeg1.1-/fn1a+* cells likely represent regenerative mesenchyme cells that also express *fn1a* (Fig. 5g). In the uninjured IOM ligament we observed almost no *mpeg1.1+* cells or *fn1a* expression. One of the most selective markers of *mpeg1.1+* cells was *lgmn*, and in situ hybridization revealed abundant *mpeg1.1+/lgmn+* cells at 1 dplt but not in the uninjured IOM ligament (Fig. 5c; Table S3). These data show that ligament transection induces a rapid recruitment of neutrophils that is followed by recruitment of *lgmn+/fn1a+* macrophages to the injury site.

### Legumain is required for efficient ligament regeneration

Given the high expression of *lgmn* in *mpeg1.1+* immune cells that home to the ligament injury site, we tested the requirement for *lgmn* in ligament regeneration by using a premature stop allele (*lgmn^sa22350^* (Fig. 6a). To characterize ECM changes during ligament regeneration, we performed Acid Fuchsin Orange G (AFOG) staining in which bone and ligament ECM stains red and cartilage and mesenchymal ECM stains blue (Fig. 6b). In wild types, we observed disappearance of the normal red ligament ECM by 3 dplt, which then began to reappear by 7 and 14 dplt and was fully restored by 42 dplt. In homozygous *lgmn^sa22350^* zebrafish (*lgmn* mutants), ligament tissue adjacent to the transection site retained ligament ECM and was not resorbed at 3 dplt as in wild types (black arrow, Fig. 6b). By 14 dplt, we observed a striking delay in ligament regeneration compared to wild types, with *lgmn* mutants displaying disorganized non-ligament ECM throughout the injury domain. At 42 dplt, *lgmn* mutants displayed reduced and highly dysmorphic regenerated ligaments consisting of several distinct nodules that failed to align along the direction of force.

**Figure 6.**
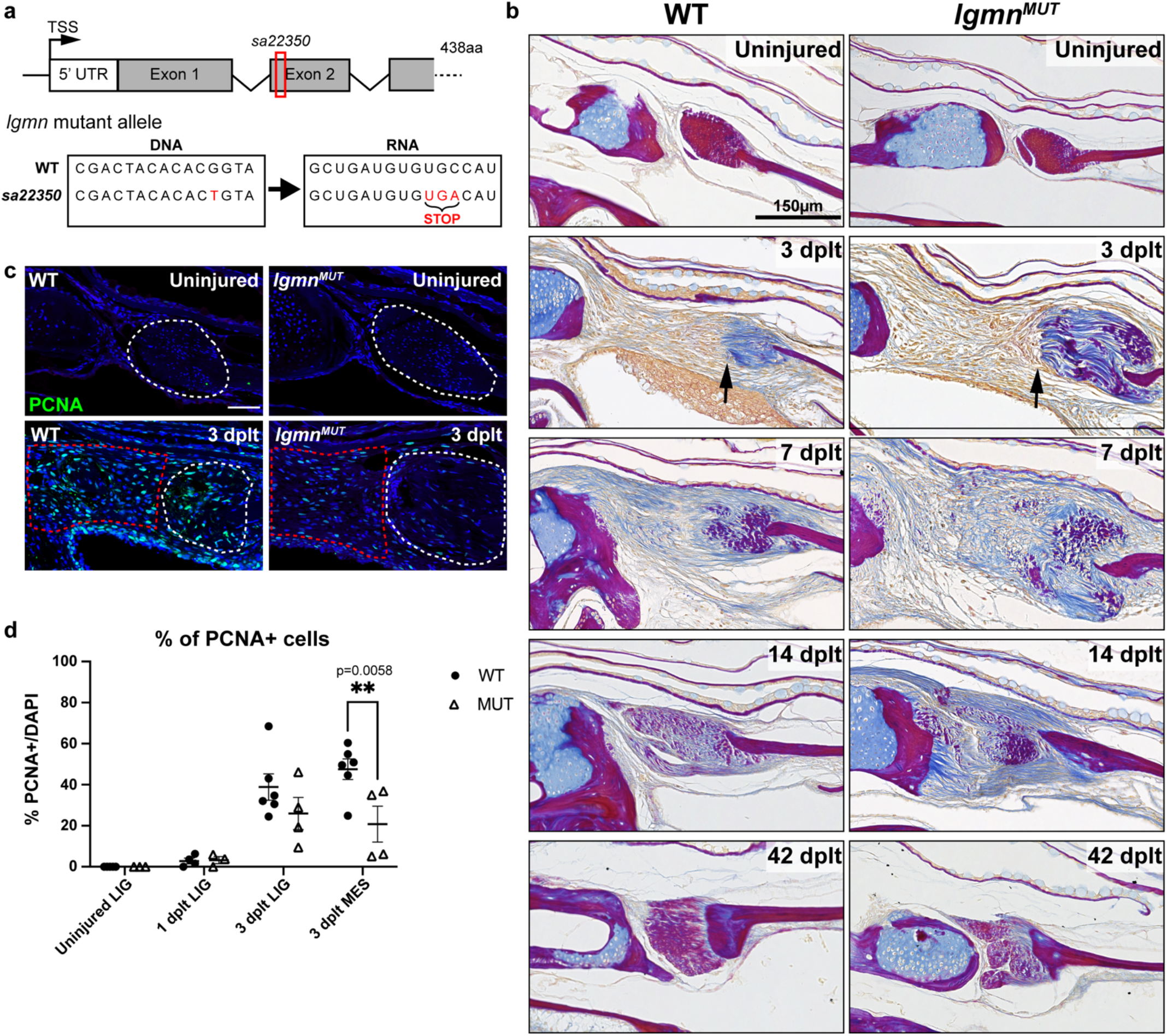
Dysregulated ECM remodeling and decreased proliferation in *lgmn* mutant after ligament transection. **(a)** Schematic of *lgmn* mutant locus showing a point mutation in exon 2 that encodes for an early stop codon. (**b)** Acid Fuchsin Orange-G (AFOG) histology staining of ECM remodeling in WT and *lgmn* mutant IOM ligament regeneration in uninjured, 3 dplt, 7 dplt, 14 dplt, and 42 dplt. Black arrow denotes interface of injured ligament bundle with invading cells. Collagen is stained blue, protein deposits are stained red/purple, and cellular cytoplasm is stained orange (n=3-5 per time point). (**c)** Representative images of PCNA immunofluorescence in WT and *lgmn* mutants in uninjured and 3 dplt IOM ligaments. Region of interest for quantification is outlined in white dashed lines for the ligament and red dashed lines for regenerative mesenchyme domain. (**d)** Quantification of percentage of PCNA positive cells normalized with total DAPI-stained nuclei (n=3-6 per time point); error bar represents SEM, p=0.005. Scale bars= 150 μm (b), 50 μm (c).

To examine the cellular bases of aberrant ligament regeneration in *lgmn* mutants, we first measured proliferation using immunofluorescence for PCNA. At 3 dplt, we observed a 27% decrease in proliferation within the regenerative mesenchyme domain of *lgmn* mutants (p=0.0058) and a trend toward lower proliferation in ligament tissue adjacent to the transection site (p=0.3369) (Fig. 6d). We also assessed potential changes in ECM gene expression in mutants (Fig. 7). In the uninjured IOM ligament of adult wild types and *lgmn* mutants, we detected very low expression of *col12a1a* and *fn1a* (Fig. 7a). At 1 dplt, we observed upregulation of *col12a1a* and *fn1a* expression in wildtype regenerative mesenchyme and ligament tissue adjacent to the transection site, with *col12a1a* but not *fn1a* showing decreased upregulation in *lgmn* mutants (Fig. 7b). By 3 dplt, *col12a1a* and *fn1a* expression had come back down in regenerative mesenchyme of wild types yet was increased in *lgmn* mutants (Fig. 7c-d). In *lgmn* mutants at 3 dplt, the majority of *fn1a+* cells were positive for the mesenchyme marker *col12a1a*, but we also observed *fn1a+/col12a1a*-negative cells likely representing invading macrophages. Reduced mesenchymal proliferation and aberrant ECM gene expression are therefore associated with the failure to properly regenerate the IOM ligament in *lgmn* mutants.

**Figure 7.**
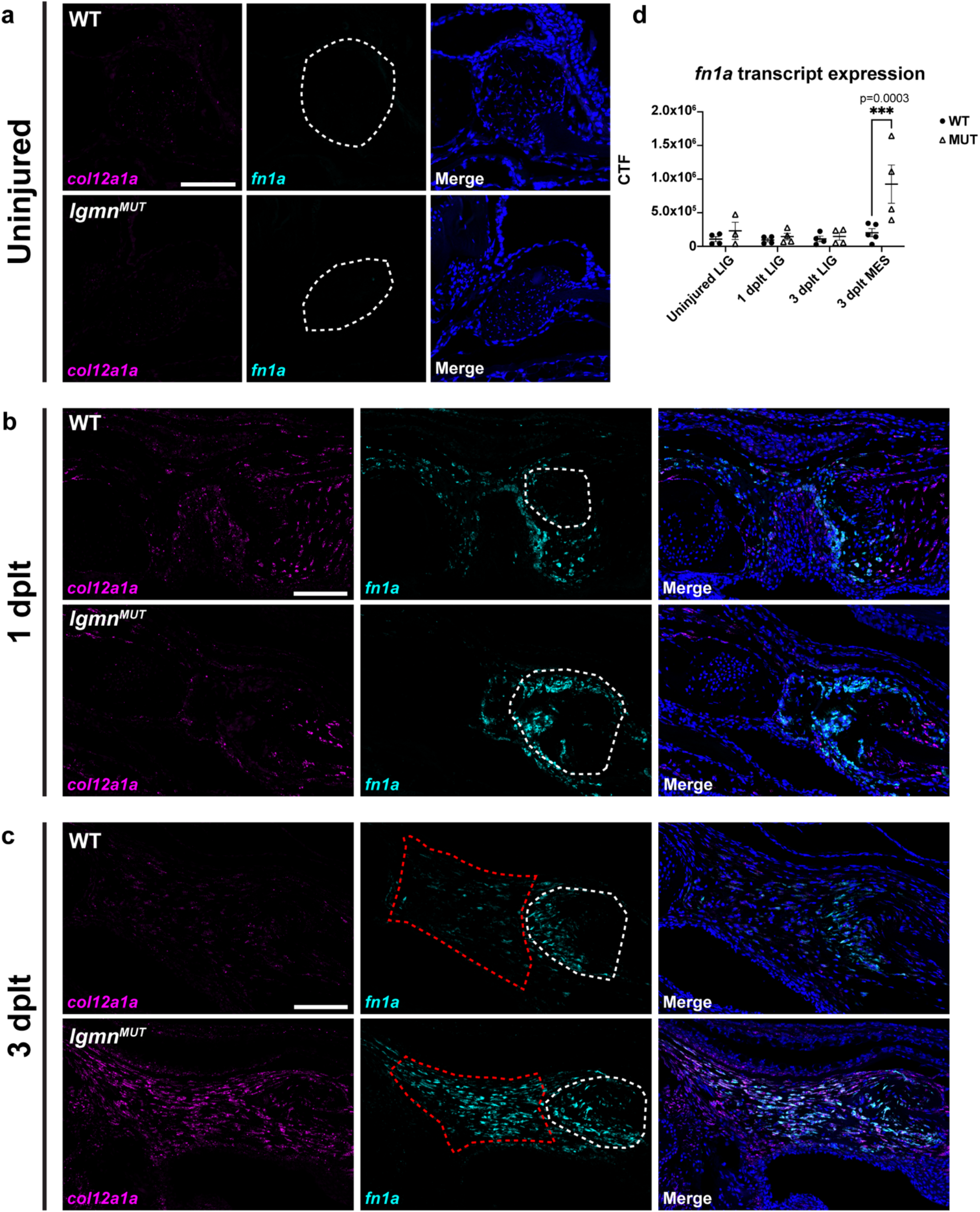
Abnormal expression of ECM factors in *lgmn* mutants. (**a-c)** Representative images of smFISH staining using *fn1a* (cyan) and *col12a1a* (magenta) probes in IOM ligaments in WT and *lgmn* mutants from uninjured (**a**), 1 dplt (**b**), and 3 dplt (**c**) (scale bar: 100μm). (**d)** Quantification of smFISH *fn1a* expression within the IOM ligament and regenerating mesenchyme. Region of interest for quantification is outlined in white dashed lines for the ligament and red dashed lines for regenerative mesenchyme (n=3-5 per time point); error bar represents SEM, p=0.0003. Scale bars= 100 μm (a-c).

## Discussion

Here we show that the IOM ligament supporting the jaw joint of adult zebrafish can fully regenerate in a month following complete transection. Lineage tracing shows that regeneration is mediated in part by the dedifferentiation, proliferative expansion, and redifferentiation of mature ligamentocytes flanking the injury site. Single-cell expression profiling further reveals dynamic changes in ECM composition during the regenerative process, mediated by both neural crest-derived mesenchyme and immune cells that home to the injury site. In particular, we identify Legumain as a key macrophage-secreted protease required for ECM remodeling, mesenchymal proliferation, and proper ligament regeneration (Fig. 8). These findings provide a paradigm for how cross-talk between the immune system and mesenchyme can promote scar-free regeneration of adult connective tissue.

**Figure 8.**
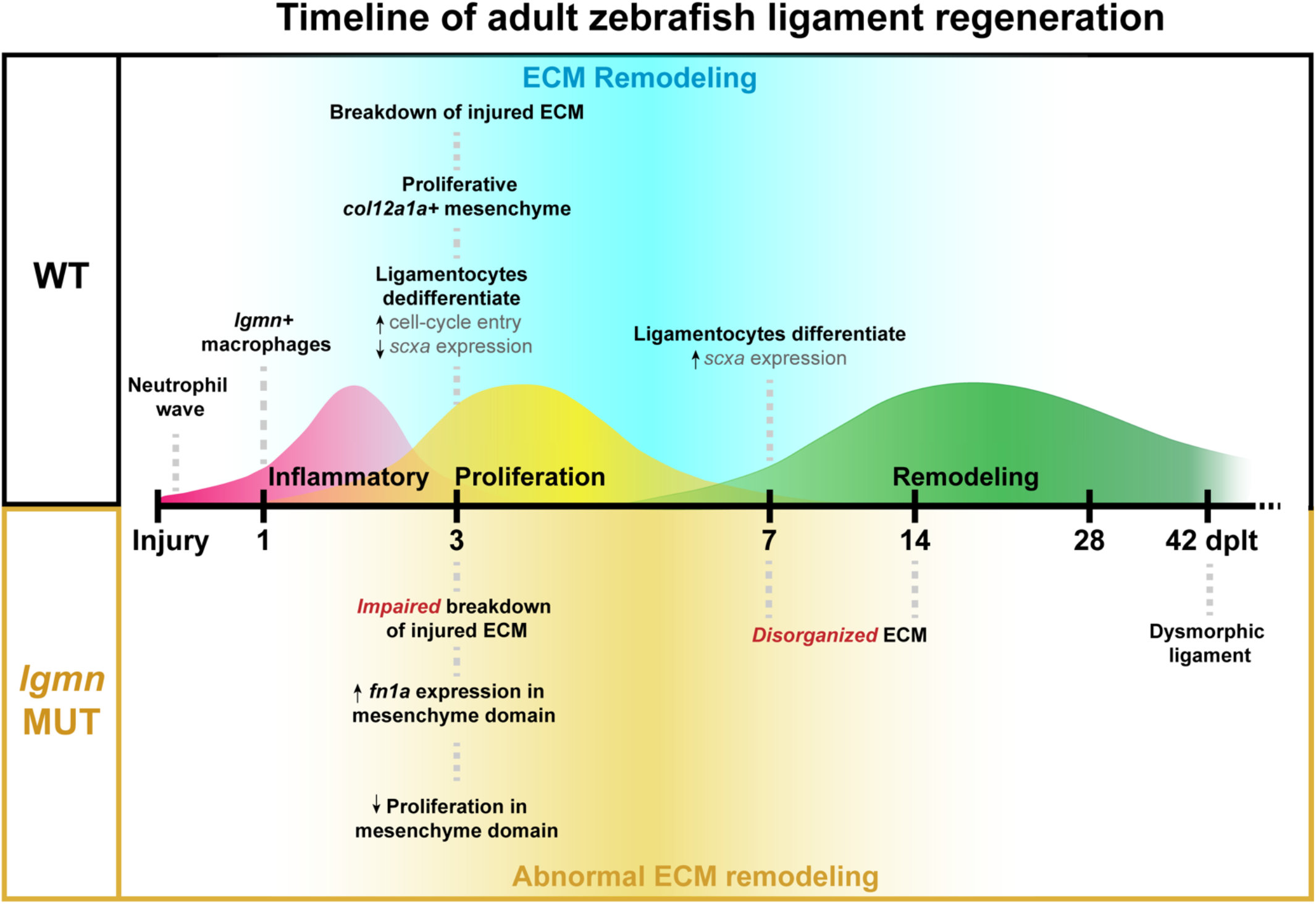
Adult zebrafish ligament regeneration timeline and major milestones. Schematic timeline of ligament regeneration in WT and *lgmn* mutants detailing the key events that occur during the three phases of healing.

Zebrafish has emerged as a high-throughput model to understand the cellular and molecular mechanisms underlying the regeneration of diverse adult vertebrate organs^13,15,26^. Our new model of adult ligament regeneration in zebrafish differs from mammalian connective tissue regeneration models, such as the neonatal Achilles tendon regeneration model in which tenocytes have retained proliferative potential from development^8^, or the periodontal ligament regeneration model that relies on a tooth-associated resident stem cell population^27^. In our model, we show that post-mitotic ligamentocytes in the adult jaw can dedifferentiate and re-enter the cell cycle after injury to fully regenerate the ligament. The concept of mature cells undergoing dedifferentiation and proliferation appears to be a common theme for adult zebrafish regeneration, for example as seen for cardiomyoctes during heart regeneration^28,29^ and osteoblasts during fin bone regeneration^30^. However, as in the fin where de novo osteoblasts also contribute to bone regeneration^31^, we cannot rule out that resident stem/progenitor cells may also contribute to ligament regeneration.

Our single-cell expression profiling reveals key changes in ECM composition in early stages of ligament regeneration. Coincident with loss of mature markers of ligamentocytes within ligament tissue adjacent to the transection site, histology reveals a rapid loss of the characteristic ligament ECM in a few days after injury. An intriguing possibility is that initial ECM remodeling facilitates the release and dedifferentiation of ligamentocytes to regenerate the ligament. At the same time, the regenerative mesenchyme bridge transiently acquires a distinct ECM signature including high levels of Type XII Collagen and Fibronectin 1, which is subsequently lost as ligamentocytes differentiate anew. These regenerative mesenchymal cells are also transcriptionally distinct from the αSMA-secreting myofibroblasts which comprise mammalian scar tissue^32,33^. Interestingly, fibroblasts expressing Type XII Collagen and Fibronectin 1 have also been implicated in heart and spinal cord regeneration in zebrafish, suggesting that a common ECM signature may facilitate scar-free repair across diverse types of organs^34,35^.

A key modulator of regeneration is the immune system. Neutrophils and macrophages have been shown to be essential for fin, spinal cord, and heart regeneration in zebrafish^36–39^. Here we used single-cell expression profiling and in vivo imaging to reveal successive waves of neutrophil and macrophage recruitment to the injury site, as well as injury-induced transcriptional changes in macrophages that accompany ligament regeneration. Macrophages strongly upregulate Fibronectin 1 expression during early phases of ligament repair, and an analogous role for macrophages in collagen deposition has been described for heart regeneration^37^. Thus, mesenchymal and immune cells may cooperate to build an ECM environment conducive to regeneration.

We found *lgmn* to be strongly expressed in macrophages recruited to the injured ligament, with loss of *lgmn* preventing proper ligament regeneration. Whereas Legumain has been shown to be critical for the resolution of fibrosis after murine myocardial infarction and obstructive nephropathy^40,41^, its role in regeneration had not been previously explored. In zebrafish *lgmn* mutants, we observed a delay in resorbing ligament tissue immediately adjacent to the injury site and a subsequent failure to regenerate an integrated and properly aligned ligament. These histological defects were accompanied by decreased mesenchymal proliferation and a failure to downregulate *col12a1a* and *fn1a* expression during later phases of regeneration. The targets of the Legumain protease in our system remain unknown. Legumain is known to cleave and activate Mmp-2, which belongs to the family of metalloproteases known to break down collagen ECM^20^. One possibility then is that Legumain may function to initiate resorption of adjacent ligament tissue, thus releasing ligamentocytes to participate in the repair process. Legumain has also been shown to degrade fibronectin ^42,43^, and thus could function later to break down the transient fibronectin-rich ECM of the regenerative mesenchyme bridge, in combination with negative regulation of *fn1a* expression. It is also possible that Legumain works in a protease-independent manner through activation of integrin αvβ3^44,45^ or αvβ1^46^. Future work will be needed to determine the relationship of ECM remodeling to transcriptional and proliferative changes of ligamentocytes and mesenchymal cells. It will also be interesting to assess potential roles of Legumain on immune cells themselves, as well as the extent to which Legumain is required in macrophages versus other cell types. In the future, a comparison of scar-free ligament regeneration in zebrafish with imperfect healing in mammals may reveal new cellular and molecular targets for therapies aimed at alleviating the substantial clinical and societal burden posed by ligament injuries.

## Materials and Methods

### Zebrafish lines

The Institutional Animal Care and Use Committees of Columbia University and the University of Southern California approved all zebrafish experiments. Published Zebrafish lines used in this study include *AB, Tübingen, lgmn^sa22350^* ^47^, *Tg(scxa:mCherry)*^52^, *Tg(actab2:loxP-BFP-STOP-loxP-dsRed)sd27*^48^, *Tg(Mmu.Sox10-Mmu.Fos:Cre)zf384*^49^, *Tg(mpx:mCherry)uwm7*^51^, and *Tg(mpeg1:EGFP)gl22*^50^. Embryos were raised in a methylene blue salt solution at 28.5°C. Juvenile and adult fish were housed in groups of 15-30 individuals. All surgical injuries were performed in size-matched 3-6 months post fertilization (mpf) adult zebrafish using standard body length (SL) measurements.

*lgmn^sa22350^* mutant allele containing a G>T point mutation resulting in a premature stop codon was used to generate homozygous *lgmn* mutants (referred to as *lgmn^MUT^*). Zebrafish were genotyped by PCR with primers flanking intron 1 (*lgmn* forward primer: AGATCTTATGATCCCAGATCCAATGACT) and end of exon 2 (*lgmn* reverse primer: GGTGAGAAAATGAAACCCGAAACTAGTCT). The 500bp PCR product was digested with NspI (New England Biolabs, R0602L) that recognizes a RCATG/Y cut site only present in mutants. Mutant alleles were then identified by presence of the cleaved 358bp and 142bp bands.

### Generation of *thbs4a_p1* transgenic zebrafish lines

The *Tg(thbs4a_p1:CreERT2)* and *Tg(thbs4a_p1:eGFP)* alleles were generated using Gateway cloning and Tol2 transgenesis. p5E plasmid including the 635bp *thbs4a_p1* enhancer and minimal *E1B* promoter sequence was generated using synthesized gBlock DNA (IDT) in a BP cloning reaction (see Supplementary File 1 for *thbs4a_p1* gBlock sequence). The *thbs4a_p1*:eGFP transgene was generated by combining *p5E-thbs4a_p1:E1B*, pME-eGFP, p3E-polyA, and pDestTol2AB2. The *thbs4a_p1*:CreERT2 transgene was generated by combining p5E-*thbs4a_p1:E1B*, pME-CreERT2, p3E-polyA, and pDestTol2CG2. Transgenes were injected into one cell stage zebrafish embryos at 30ng/ul plasmid DNA with 50ng/ul Tol2 mRNA. Injected F0 animals were raised to adulthood and outcrossed to *Tübingen* or *Tg(actab2:loxP-BFP-STOP-loxP-dsRed)sd27* animals and F1 founders were screened for the presence of eye CFP (*thbs4a_p1*:eGFP) or heart GFP and successful 4-OHT conversion in progeny (*thbs4a_p1*:CreERT2), respectively.

### Adult ligament transection surgery

Interopercular-mandibular (IOM) ligament transection surgery was adapted from Smeeton et al, 2022^18^. Briefly, adult zebrafish were anesthetized with Tricaine MS-222 and restrained in a damp sponge. Using 3-mm Vannas spring scissors (Fine Science Tools, cat. #1500000), the IOM ligament was severed with a single cut and a pull test performed on the IOP bone to confirm transection. Fish were then revived in clean system water, housed on-system, and received daily post-operative health checks for three days.

### Histology, in situ hybridization, and immunohistochemistry

Zebrafish were processed for paraffin embedding and histological sectioning as previously described (Askary et al, 2016). Paraffin sectioning was performed using Thermo HM355S automatic microtome.

For AFOG staining, 5uM paraffin sections were deparaffinized with Hemo-De and re-hydrated through an ethanol series (100, 90, 70, 50, 0) diluted with water. Sections were then fixed in 4% PFA for 5 min at RT, then washed 3x for 5min with 1xPBS and then washed 1x for 5 min in water. Following washes, samples were incubated in Bouin’s solution (Sigma-Aldrich, HT10132) that was preheated to 60°C for 2-hours and cooled to room temperature for another hour. Slides were then washed 5-6x for 5 min with water and then rinsed in 1% Phosphomolybdic acid (Sigma-Aldrich, HT153) for 10 min and rinsed in water for 5 min. Slides were then incubated in AFOG staining solution (0.5% aniline blue, 1% orange-G, and 1.5% acid fuschin, pH 1.09; Sigma-Aldrich, 415049, O3756, F8129) for 7min at RT and then rinsed 1x for 2min with water. Samples were dehydrated with a series of ethanol dilutions (75, 90, 100%) and 1x in Hemo-De for 5min before mounted with Cytoseal (Epredia, 23-244256).

For Toluidine Blue/Fast Green staining, zebrafish tissue was fixed and paraffin embedded according to the aforementioned paraffin embedding protocol. Paraffin blocks were sectioned using HM 355S automated microtome and MX35 microtome blades at 5μm. Sections were floated in DEPC-treated water (Sigma-Aldrich, D5758) in a waterbath and collected onto Leica adhesive slides, then allowed to air dry at room temperature at least overnight. Slides were then processed for toluidine blue staining with the following procedure: two 5-minute washes in xylene substitute Hemo-De, two 1-minute washes in 100% ethanol; serial washes (90%, 70%, and 50%) in EtOH/H2O for 1 minute per wash; and three 1-minute washes in tap water. Slides were then placed in 0.04% toluidine blue solution (0.08g toluidine blue in 200mL of 0.1M sodium acetate, pH 4.0; Sigma-Aldrich, 89640) for 10 minutes, followed by three 1-minute washes in tap water. Slides were then placed in 0.1% fast green solution (0.2g fast green in 200mL water; Fisher Scientific, BP123-10) for 3 minutes, followed by three 1-minute washes in tap water. Stained slides were then given two 1-minute washes in 100% isopropanol and two 1-minutes washes in Hemo-De. Lastly, coverslips were mounted on the slides using CytoSeal and allowed to air dry for 30 minutes to overnight.

### Tissue clearing

CUBIC tissue clearing protocol was adapted from (Susaki et al., 2014) for use with zebrafish^53^. Whole adult zebrafish were fixed overnight in the dark at 4°C in 4% PFA. Tissue was then washed twice in PBS for at least 30 minutes per wash and heads were separated from the trunk of the body. At a ratio of 5 zebrafish heads per 50mL of solution, tissue was incubated in the dark at 37°C in CUBIC-1 solution with gentle rocking for at least 3 days. Tissue was then transferred to new CUBIC-1 to incubate until tissue was transparent and iridophores were cleared, 2-4 days. Tissue was then washed twice in PBS for at least 30 minutes per wash, then immersed in 20% sucrose in PBS at room temperature until tissue was no longer floating in solution and once again opaque, 2 hours to overnight. Lastly, the tissue was immersed in CUBIC-2 solution and incubated in the dark at room temperature with gentle rocking for at least 1 day before imaging. Fluorescent microscopy images were captured using a Leica SP8 confocal microscope and processed in Fiji. CUBIC-1 was prepared as a mixture of 25 wt% urea, 25 wt% N,N,N’,N’-tetrakis(2-hydroxypropyl) ethylenediamine (Fisher Scientific, AAL16280AP) and 15 wt% Triton X-100. To minimize water evaporation, solution was prepared with heat not exceeding 100°C and inside bottles with caps just screwed on. Deionized water was heated prior to the addition of urea. Once urea was dissolved and the solution was no longer cold, N,N,N’,N’-Tetrakis(2-hydroxypropyl)ethylenediamine was mixed until homogenous. Solution was removed from heat and allowed to cool to room temperature, then Triton-X-100 was added with gentle stirring to minimize the production of bubbles. CUBIC-1 solution was stored in the dark at room temperature and used within 1 week.

CUBIC-2 was prepared as a mixture of 50 wt% sucrose, 25 wt% urea, 10 wt% 2,2’,2”-nitrilotriethanol (Fisher Scientific, AAL044860E), and 0.1% (v/v) Triton X-100. To minimize water evaporation, solution was prepared with heat not exceeding 100°C and inside bottles with caps just screwed on. Deionized water was heated prior to the addition of urea. Once urea was dissolved and the solution was no longer cold, 2,2’,2’’-Nitrilotriethanol was mixed until homogenous. Sucrose was mixed with continued heat until the solution was clear with a pale yellow tint. Solution was removed from heat and allowed to cool to room temperature, then Triton-X-100 was added with gentle stirring to minimize the production of bubbles. CUBIC-2 solution was stored in the dark at room temperature and used within 1 week.

### Immunofluorescence and smFISH

We collected 5μm formalin-fixed paraffin-embedded (FFPE) tissue sections. For immunofluorescence, sections were deparaffinized with Hemo-De and rehydrated through an ethanol series, then subjected to antigen retrieval in sodium citrate buffer, blocked with Agilent Dako protein block (cat. X090930-2), and incubated with primary antibody (PCNA, 1:1000; Sigma-Aldrich, P8825) overnight at 4°C, washed, and incubated with secondary antibody (1:500; Invitrogen, PIA32727). Slides were mounted and counterstained with Fluoromount-G + DAPI (Southern Biotech, 0100-20). Single-molecule fluorescence *in situ* hybridization was performed on 5μm FFPE sections according to manufacturer guidelines using RNAScope Multiplex Fluorescent Reagent Kit v2 Assay (ACD Bio). Slides were treated with 1x Target Retrieval Reagent for 4min in a steamer. RNAScope probes used were Dr-acta2 (508581-C4), Dr-col12a1a (556481-C2), Dr-DsRed-C2 (custom), Dr-fn1a (1097911-C1), Dr-scxa (564451), Dr-lgmn (1003381-C2), Dr-mpeg1.1 (536171-C3), and Dr-thbs4a (812151) from ACD Bio. Opal 520, 570, and 690 fluorophores were used to visualize expression (1:1,000; Akoya Biosciences, cat. #FP1487001KT, #FP1488001KT, and #FP1497001KT). Sections were counterstained and mounted with Fluoromount-G + DAPI.

### Drug treatments

Adult *thbs4a_p1*:CreERT2 zebrafish lines were converted with three 5μM (Z)-4-Hydroxytamoxifen, ≥98% Z isomer (Sigma-Aldrich, H7904) overnight treatments in the dark. Following drug treatment, adults were screened for conversion using Leica M165FC stereo microscope and then washed 2 × 1 hour in system water before being re-housed on-system.

### Single cell RNA sequencing

Ligament transection (n=22 joints per 1 dplt and 3 dplt timepoints), sham surgery (n=22 joints per 1d and 3d timepoint) was performed on size-matched *Sox10Cre;actb2:loxP-BFP-loxP-DsRed* fish at 3-4 mpf. The jaw joint region of each fish and uninjured control (25 joints) was microdissected and incubated in protease solution [0.25% trypsin (Life Technologies, 15090–046), 1 mM EDTA, and 400 mg/ml collagenase D (Sigma-Aldrich, 11088882001) in PBS] with mechanical dissociation by nutation and agitation with a P1000 pipette every 5 minutes for 1hr at 28°C. Cells were FACS sorted and immediately processed to create single cell RNA-Sequencing libraries using the Chromium Single Cell 3’ Library & Gel Bead Kit v2 (10X Genomics), according to the manufacturer’s recommendations. To determine cell count, libraries were sequenced at the CHLA Center for Personalized Medicine Genomics Core using a MiSeq Nano v2 (Illumina, San Diego, CA) with 26bp sequencing for Read1, 8bp sequencing for Index1, and 120bp sequencing for Read2. Libraries were pooled for even coverage per cell and were sequenced at the CHLA Center for Personalized Medicine Molecular Genomics Core on the NextSeq 550 (Illumina, San Diego, CA) with 26bp sequencing for Read1, 8bp sequencing for Index1, and 120bp sequencing for Read2. All samples were sequenced to an average read depth of greater than 130,000 reads per cell. CellRanger v3.0.0 (10X Genomics) was used with default parameters for alignment to GRCz11 to generate gene-by-cell count matrices.

Analyses of scRNAseq libraries were performed using R version 4.0.5 in RStudio with Seurat version 4.0.2. CellRanger output for all samples were converted to Seurat objects and merged, then filtered for cells with between 400 and 1,900 unique features per cell, with under 5% mitochondrial RNA detected per cell to remove low-quality cells, doublets, or dying cells. Data were log-normalized before selecting 2,000 variable features, then scaled using all features expressed in at least 3 cells. Linear dimensional reduction was then performed using principal component analysis to compute 80 principal components. Statistical significance for each principal component were determined with a jackstraw permutation, which identified 68 significant principal components (p < 0.05). Principal components 1-68 were used for K-nearest-neighbor graph construction (KNN). Clustering was performed at low-resolution (r = 0.02) to identify broad superclusters for downstream analysis (Fig. 2a, 2b). Clusters and gene expression were graphed using UMAP-non-linear dimensional reduction to preserve global variability.

Analysis of scRNAseq data from neural-crest-derived cells were performed by subsetting and merging the superclusters containing cells from the DsRed-enriched samples (Fig. 3c). Using the unprocessed data from these cells, gene expression data were again processed with lognormalization, feature selection, and feature scaling. Linear dimensional reduction was then performed using principal component analysis to identify 50 principal components, of which 38 were deemed significant using a jackstraw (p < 0.05). Principal components 1-38 were used for KNN and graphed using UMAP-non-linear dimensional reduction before clustering at r = 0.95. Cluster 13, which showed uncharacteristically high unique feature counts and markers for epithelial cells (*epcam*, *cdh1*), were excluded as a likely doublet cluster. Markers for each cluster were identified with function FindAllMarkers using a Wilcoxon Rank Sum test for features expressed in at least 10% of the cluster and a minimum of 0.2 log-fold-change higher expression than the other cells in the neural crest-derived superclusters.

Analysis of scRNAseq data from immune cells were performed by subsetting and merging the superclusters Immune 1 and Immune 2 (Fig. 3b). Using the unprocessed data from these cells, gene expression data were again processed with log-normalization, feature selection, and feature scaling. Principal components 1-50 were used for KNN and graphed using UMAP-non-linear dimensional reduction before clustering at r = 0.9. Markers for each cluster were identified with the function FindAllMarkers using a Wilcoxon Rank Sum test for features expressed in at least 10% of the cluster and a minimum of 0.25 log-fold-change higher expression than the other cells in the immune superclusters.

### Imaging

Images of histological sections were captured using a Keyence BZ-X810 microscope and software, then processed in Fiji. Live imaging was performed using a Leica M165FC stereo microscope with Leica K5 sCMOS camera and LASX software v3.7.4.23463. Fluorescence imaging was performed using a Leica SP8 confocal microscope with LASX software v3.5.7.23225.

### Quantification and statistical analysis

Proliferation was quantified as a percentage PCNA+ cells of total DAPI+ cells in a region of interest (ROI) in the uninjured ligament domain, ligament stub domain, or regenerative mesenchyme domain and scored by an observer blinded to treatment and/or genotype. WT and *lgmn* mutant PCNA staining and quantification were performed concurrently and WT quantification is shown both in Fig. 2f and as WT controls in Fig. 6d. ImageJ was used to quantify the smFISH images. Ligament domains and mesenchyme domains were drawn based on regional nuclear morphology in the tissue section. To calculate the corrected total fluorescence, an ROI was drawn to measure the integrated density for each channel using the same ROI for each channel. To account for tissue autofluorescence and background signal, 3 background regions of each image were measured, and the average mean gray value was calculated. To calculate the corrected total fluorescence (CTF), we used the following formula CTF = Integrated Density-(area of ROI*background mean).

All statistical analysis was carried out using GraphPad Prism 9. Proliferation in WT animals was performed by ordinary 1-way ANOVA with Tukey’s multiple comparison test. Comparison of WT to *lgmn* MUT proliferation was performed by 2-way ANOVA with Šídák’s multiple comparisons test.

## Data availability

Raw and processed scRNA sequencing files have been deposited in GEO and can be retrieved using accession number GSE224197.

## Acknowledgments

We thank Joshua Barber, Megan Matsutani and Jennifer DeKoeyer Crump for fish care, Jeffrey Boyd at the USC Flow Core, and Dave Ruble at the CHLA Sequencing Core. The authors declare no conflicts of interest.

## Author Contributions

J.S. devised the surgical injury model. J.G.C. and J.S. conceived the project. T.A., J.M., E.G., D.S., M.B., E.C., G.L., and J.S. performed the injury experiments and analyses. K.-C.T., P.F., and J.S. performed the scRNAseq and pre-processed the data. J.S. generated and screened the *thbs4a_p1*:eGFP zebrafish line. J.G.C., H.-J.C., and J.S. generated and screened the *thbs4a_p1*: CreER zebrafish line. T.A. and J.S. performed the scRNAseq data analysis. T.A., J.M., E.G., and J.S. wrote the first draft of the manuscript with editing by J.G.C. and input from all authors.

