## Supplemental Figures for "Ligament injury in adult zebrafish triggers ECM remodeling and cell dedifferentiation for scar-free regeneration"

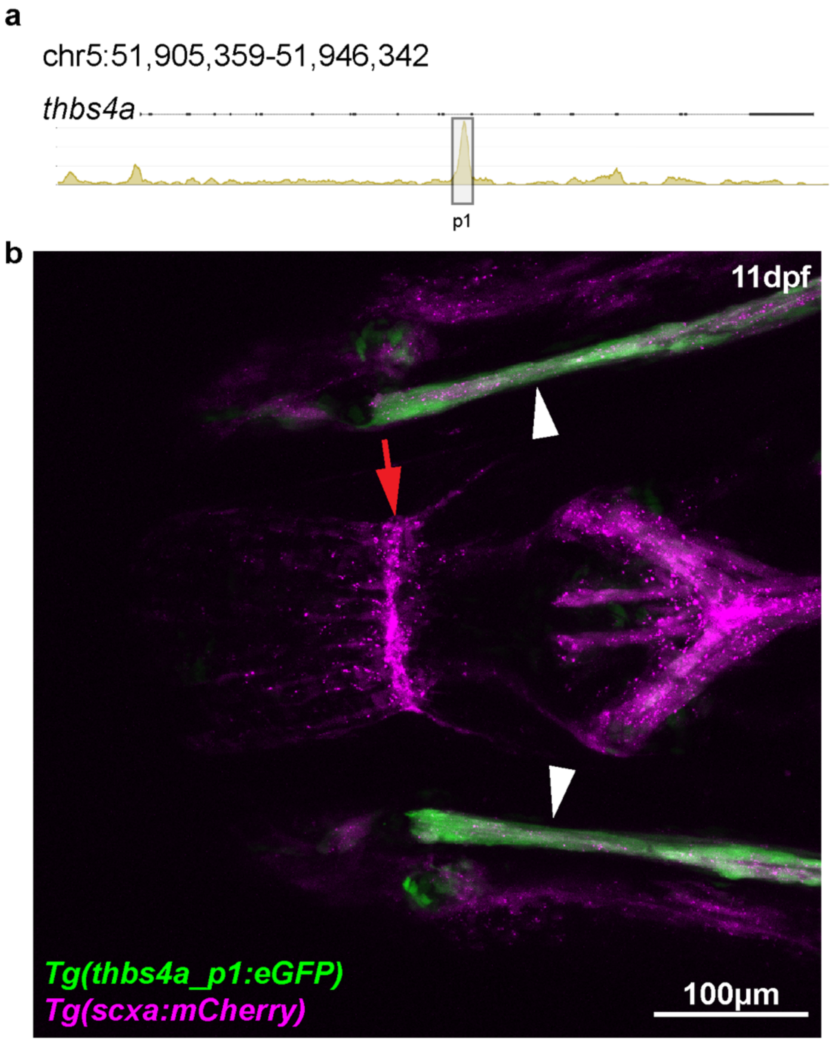


**Figure S1. Identification of *thbs4a_p1* enhancer and the developmental expression of *thbs4a_p1:eGFP* and *scxa:mCherry* in 11dpf embryos.**

(**a**) Zebrafish *thbs4a* locus showing the location of the *thbs4a_p1* enhancer sequence. (**b**) Representative images of WT transgenic *Tg(thbs4a_p1:eGFP/scxa:mCherry)* zebrafish at 11 days post fertilization (dpf). Red arrows denote the midline tendon that is single positive for *scxa:mCherry* expression. White arrowheads denote double positive *thbs4a_p1:eGFP/scxa:mCherry* signal in the bilateral IOM ligaments. *thbs4a_p1:eGFP* expression in the craniofacial region is limited to ligament cells and articular cartilage in the developing lower jaw joint (n=3). Scale bar= 100 μm.

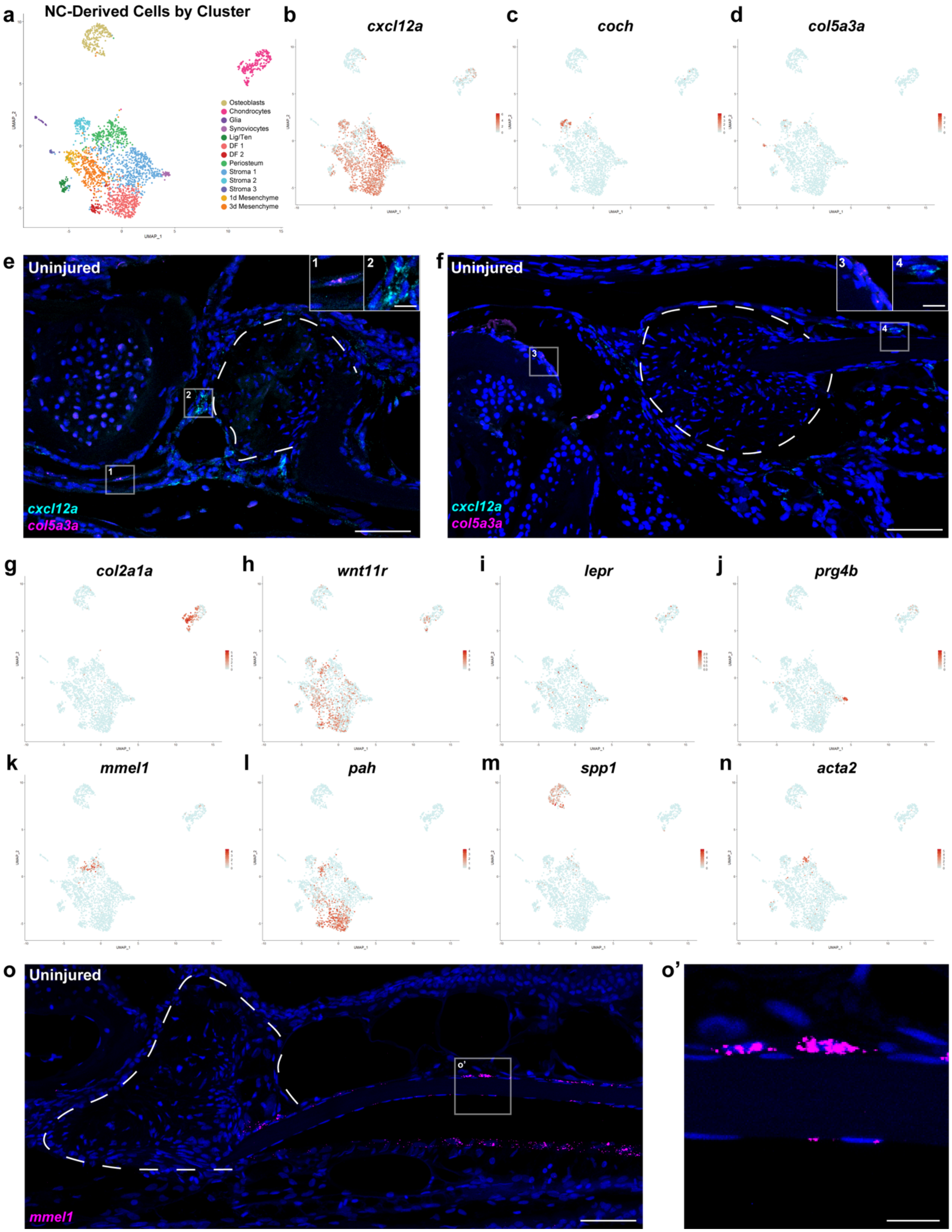

a
NC-Derived Cells by Cluster
Osteoblasts
Chondrocytes
Glia
Synoviocytes
Lig/Ten
DF 1
DF 2
Periosteum
Stroma 1
Stroma 2
Stroma 3
1d Mesenchyme
3d Mesenchyme
UMAP_1
UMAP_2
b
cxcl12a
c
coch
d
col5a3a
e
Uninjured
cxcl12a
col5a3a
f
Uninjured
cxcl12a
col5a3a
g
col2a1a
h
wnt11r
i
lepr
j
prg4b
k
mmel1
l
pah
m
spp1
n
acta2
o
Uninjured
mmel1
o'

**Figure S2. scRNAseq and smFISH characterization of tissue heterogeneity in adult zebrafish joints.**

**(a)** UMAP illustrating clustering of neural crest-derived cells in early ligament regeneration. **(b-d)** FeaturePlots representing the expression of markers for clusters Stroma 1 (*cxcl12a*), Stroma 2 (*coch*), and Stroma 3 (*col5a3*). **(e-f)** RNAscope smFISH for *cxcl12a* (cyan) and *col5a3* (magenta) near uninjured ligament (white dashed line) (n=3). **(g-j)** FeaturePlots showing the expression of published markers of deep (*col2a1a*, *wnt11r*), intermediate (*lepr*), and shallow (*prg4b*) articular chondrocytes. **(k-n)** FeaturePlots illustrating markers of subsets of the Periosteum cluster, containing subsets of periosteum (*mmel1*) resembling dermal fibroblasts (*pah*) or pre-osteoblasts (*spp1*), as well as perivascular cells (*acta2*). **(o-o')** RNAscope smFISH demonstrating *mmel1* transcripts (magenta) in periosteal cells lining the IOP bone (n=3). Scale bar = 50 µm (e,f,o), 10 µm (o').

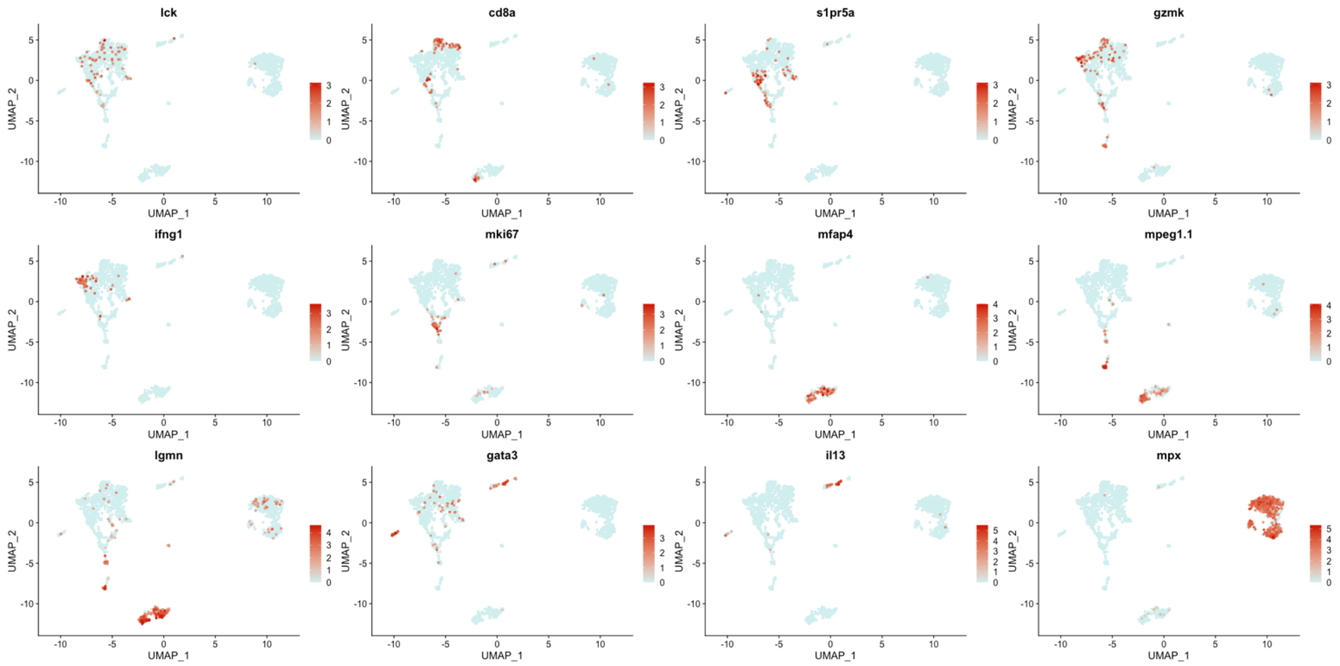


**Figure S3. scRNAseq feature plot expression for cluster-defining genes in immune populations.**

Feature plots of mRNA expression of immune cluster marker genes.

**Supplementary File 1**

*thbs4a_p1* sequence
> NC_007116.7:51926588-51927222 Danio rerio strain Tuebingen chromosome 5, GRCz11 Primary Assembly

CAAATATATATAGAGGTATACTGTCTAAAATGTCACTGAATGGAGCAAACTCCATCAGCAGA
AAAGCGTTGCACTGTTTTAGACTTGCATACAGTGACATCTTTGTGTGCGGTTATGTCACAAG
GTGGCAGTGTTTCAGAAAAAAAGACTTCTGTGGAATAGTTTCCACATGCCAGCAGCTTTTGG
CTTGGAATCTCTGTGGAAATGTGGTCGCTCGAAGGAGCCATGACCAAGCTTTAAAGCATCTA
ATCAAGTGTCTGGATTTCTGCCTTGATGAGCAGACCCTAAATGTCAAACAACATCTGGACTC
GATTTTACACATTTCAGACTGGAAGGAATAATTGTGCATTTAAAAGAAAATGATCAGTGTGG
TGCAGTAAATGCAGCTAATATCTTTACAGGTCACTTCAGTTTTACATGACCCGCGCAAACTA
AAGAGCGCTCTAATTTGAGCTTTTAAAACACAGTAGCTTCTGTTTTTCCTCATAAAAATGACC
TAAGCTCTGGAAAATCCAGATAGAAAACAGGCCCGAAGGGCAAAAAGTATATTTAAAACAC
ATCACCCCATTTCTATGAATAAAAATGTGTCTTTGGCCACTTAGCAGCCGGATGACACGTGA
AATGTTACTGCAGCA

**Supplemental Videos S1-4.** 3D rendering of tissue-cleared imaging of *thbs4a_p1*:eGFP in uninjured (Video S1), 1 dplt (Video S2), 7 dplt (Video S3), and 28 dplt (Video S4) IOM ligaments.

**Table S1. scRNAseq cell counts by sample**

**Table S2. Marker genes of neural crest lineage scRNAseq clusters**

**Table S3. Marker genes of immune scRNAseq clusters**
